# High diversity of type I polyketide genes in *Bacidia rubella* as revealed by the comparative analysis of 23 lichen fungal genomes

**DOI:** 10.1101/2022.02.10.479404

**Authors:** Julia Gerasimova, Andreas Beck, Silke Werth, Philipp Resl

## Abstract

Fungi involved in lichen symbioses produce a large array of different secondary metabolites. The high diversity of those substances has been known for decades and are often considered characteristic for taxonomical delimitation in lichen-forming fungi. Polyketides, the most common secondary metabolites, are synthesized by the Type I Polyketide synthases (TI-PKS) comprised of different enzymatic domains. We present a phylogenetic overview of Type I PKS genes recovered from the *de-novo* sequenced genome of *Bacidia rubella* in the context of additional twenty-one fungal genomes from the largest radiation of lichen-forming Ascomycetes (Lecanoromycetes) as well as the lichen-forming Eurotiomycete, *Endocarpon pusillum*. Using *de-novo* gene prediction and functional annotation combined with a phylogenetic analysis, we provide insights into the biosynthetic potential and PKS gene diversity of lichen-forming fungi. We discuss genes predicted in the lichen-forming fungal genomes in relation to previously characterized PKS genes from other fungi and bacteria. Our results reveal a high number of biosynthetic gene clusters and their gene domain composition. PKS gene content outnumbers known secondary substances produced by the lichen-forming fungi. We were able to assign putative functions to several of those PKS genes *in silico*, based on similarity to already characterized genes. In particular, we identified a putative PKS23 gene of *Bacidia rubella*, producing the common lichen substance atranorin. However, we also found that several lichen-forming fungi still possess homologs of different biosynthetic genes without producing the corresponding substances in detectable amounts. Although many PKSs remain without functional assignments in our analysis, our findings highlight that genes from lichen-forming fungi represent an untapped source of novel polyketide compounds. However, additional experimental approaches are necessary to link biosynthetic genes and secondary metabolites with confidence.

## Introduction

Fungi synthesize an extensive array of natural products, termed secondary metabolites, with roles in defence, self-protection and development [1,2]. Similar to their functions, these metabolites are chemically diverse. Based on their properties and the core enzymes and precursors involved in their biosynthesis, four major groups can be distinguished: polyketides, non-ribosomal peptides (NRPS), terpenoids and tryptophan derivatives [3]. The genes encoding most fungal secondary metabolites are located adjacent to each other (*i.e*., “clustered”) in the genome [2,4].

In lichen forming fungi, polyketides are the most common class of secondary metabolites [5,6]. Although lichen polyketides can sometimes be produced by mycobionts grown axenically under appropriate conditions [7–10], many lichen polyketides are only formed in intact symbioses [11,12]. Lichen polyketides are synthesized by Type I polyketide synthases (TI-PKS) [13], whose closest structural and functional analogue is the mammalian fatty acid synthase [14]. In the minimal configuration, the domain structure of TI-PKSs includes a ketoacyl synthase (KS), an acyltransferase (AT), and an acyl carrier protein (ACP), which are essential for polyketide synthesis [3]. This configuration can be supplemented by additional domains, such as starter unit-ACP transacylase (SAT), ketoreductase (KR), dehydratase (DH), enoyl reductase (ER), methyltransferase (CMeT), and thioesterase (TE) [3].

The presence of optional domains has led to the classification of fungal PKSs into three subgroups based on their ability to perform redox reactions. The first subgroup comprises the non-reducing (NR) PKSs, which lack reductive domains and mainly produce aromatic polyketide [13,15]. The second subgroup contains partially-reducing (PR) PKS, typically having a single KR or KR and DH domains. Finally, reducing (R) PKSs contain a complete set of reductive domains, viz. KR, DH, and ER. All three subgroups can be found in lichen-forming fungi, and their diversity has not gone unnoticed. Previous work suggests various ecological roles of polyketides in lichen-forming fungi ranging from light-screening and chemical weathering to allelopathic effects and herbivore defence [3,6,16,17]. More broadly, secondary metabolite profiles including polyketides are often characteristic for taxonomic groups and are thus extensively used to distinguish different lichen-forming fungi.

The first PKS gene from lichen-forming fungi has been cloned and analyzed by Armaleo *et al*. [18] and in recent years, secondary metabolite research has benefited extensively from genome mining approaches in bacteria, fungi, and plants, uncovering hidden diversity (e.g., [19–21]). The fact that genes encoding natural product biosynthetic pathways are often clustered in the genome [2] facilitates the identification of biosynthetic gene clusters (BGCs) in genome sequences. Moreover, in many cases, the chemical structures of their products can be predicted to a certain extent, based on the interpretation of the biosynthetic logic of enzymes encoded in a BGC and their similarity to known counterparts [22–25]. Despite this progress, for most secondary metabolites of lichen-forming fungi, the corresponding biosynthetic genes remain uncharacterized. Observing that lichen-forming fungi contain far more biosynthetic genes than characterized chemical compounds also complicates clear gene assignments to a particular metabolite. Consequently, previous studies have focused only on a few well-known compounds, e.g., usnic acid, grayanic acid, and atranorin (e.g., [18,25–29]). However, the increasing number of available lichen-forming fungal genomes provides an untapped resource for identifying additional PKS genes [30].

Here we investigate the diversity of BGCs in the *de-novo* sequenced *Bacidia rubella* (Hoffm.) A. Massal. genome within a two-level comparative genomic framework. First, we aim to provide insight into the secondary metabolite biosynthetic potential of *B. rubella* in the context of the family Ramalinaceae by analyzing the genomes of *Bacidia gigantensis*, *Ramalina intermedia* and *R. peruviana*. The secondary metabolites of these species are known and there is overlap in the substance profiles between different species. This allows us to detect the presence and absence of BGCs of previously characterized PKS genes encoding for atranorin biosynthesis within closely related species. We then extend this approach to include another nineteen publicly available fungal genomes from the Lecanoromycetes obtained from pure cultures, with high genome completeness and little contamination. By employing an *in-silico* approach combined with phylogenetic reconstructions, we assign putative functions to BGCs of lichen-forming fungi. Our comparative genomic results are in line with previous work to indicate a high diversity of BGCs in lichen-forming fungi beyond what can be observed in their chemical profiles. This indicates a tremendous potential of the here-studied genomes to produce different secondary metabolites, with direct relevance for natural product research and production.

## Material and methods

### *In vitro cultivation of the* Bacidia rubella *mycobiont*

The lichen-forming fungus *Bacidia rubella* was axenically cultivated from a specimen collected from Germany (Bavaria, Lkr. Neuburg-Schrobenhausen, Markt Rennertshofen, south-east of Bertoldsheim, Naturwaldreservat “Mooser Schütt”; mixed forest, on bark of the trunk of *Fraxinus* sp., ca. 1.0 m above the ground, ca. 400 m asl.; M-0307710) in April 2019. The mycobiont culture was obtained from a multispore discharge of a single apothecium of *B. rubella* following the method of Yoshimura *et al*. [31]. Briefly, young and middle-aged apothecia were detached from the thallus and soaked in sterile water for about an hour. Then they were fixed to the top of a Petri dish lid using petroleum jelly while keeping the lid slightly open to let the apothecia dry slowly. We used a Bold’s Basal Media (BBM) with doubled nitrate (25 g/L) as an initial substrate. Upon germination, the spores were transferred to a malt-yeast extract medium (Lichen medium) [32]. The mycobiont cultures were stored in a growth chamber (Wachstumsschrank Binder KBWF 720; Tuttlingen, GmBH) at 16°C and 60% relative humidity. They were subcultured every two to three months until sufficient biomass for genomic analysis were obtained (ca. one year).

### DNA isolation and sequencing

To obtain concentrated high molecular-weight genomic DNA, about 1 cm^2^ of mycelium was taken and ground in liquid nitrogen with a pre-cooled pistil. Genomic DNA was isolated using the MagAttract HMW DNA Kit following the manufacturer’s protocol and subsequent purification steps with magnetic beads (HMW genomic DNA from fresh or frozen tissue protocol), resulting in a total yield of 1 μg. The final concentration was 9.53 ng/μL measured with a NanoDrop 1000 spectrophotometer (Peqlab Biotechnologie, GmbH) and Qubit 4 Fluorometer (ThermoFisher Scientific) using 1X dsDNA HS Assay Kits. DNA extraction, PCR amplification and Sanger sequencing (using nrITS primers) were performed to evaluate possible contamination and confirm the cultures’ identity. First, a paired-end library was constructed using Illumina DNA Prep (earlier known as Nextera DNA Flex Library Prep) and sequenced on NovaSeq 6000 (NovaSeq SP 150bp Paired-end Flow Cell, Illumina) at the Biomedical Sequencing Facility (BSF, Vienna, Austria). A total concentration of 0.2 μg was used for Illumina library preparation. To supplement the Illumina data, we prepared several Oxford Nanopore libraries using the SQK-LSK109 kit and sequenced them on a MinION sequencer using R9.4.1 flow cells (altogether, four runs were conducted). The total yield of DNA was in the range of 0.12-0.31 μg.

### Data generation and initial read-quality assessment

To obtain a high-quality genome, we used a hybrid assembly approach using Illumina short-reads and Oxford Nanopore long reads. In total, we produced 17 Gbps of raw paired-end Illumina reads with 500x coverage. Raw data inspection with FastQC v0.11.7 [33] indicated high-quality reads (quality score (Q) above 34) and no excessive adapter contamination or read-duplication. Basecalling of Nanopore raw signals was performed using Flappie v2.1.3 (github docker://reslp/flappie:4de542f; reference) into a total of 22 Gbps of raw sequences up to 94.5 Kb of read length.

### Genome completeness and quality assessment

The paired-end Illumina reads and unpaired Nanopore reads were assembled using hybridSPAdes [34] with k-mer sizes of 33, 55, 77 and 127. To examine the assembly for potential non-target contigs, *de-novo* assembly was subjected to BLASTX using DIAMOND [35] against a custom database comprising the protein sets of the NCBI nr database (downloaded in July 2018). The results of this DIAMOND search were used as input for blobtools v1.1.1 [36]. The final results showed the absence of foreign contigs; therefore, further filtering was unnecessary. The quality of the assembly and genome statistics were assessed using QUAST v5.0.2 [37]. The completeness of the genomes was assessed using all single-copy BUSCO genes of the ascomycota_odb9 set (part of phylociraptor pipeline, see below). An overview of BUSCO results is given in Table 4.

### Dataset construction

The dataset consists of the genomes from the *de-novo* sequenced genome of *B. rubella* and twenty-two additional representative genomes of Lecanoromycetes, the largest radiation of lichen-forming fungi. We included in our study twenty-three genomes obtained from pure fungal cultures with the exception of *B. gigantensis* and *R. intermedia*, which were obtained by sequencing the whole thallus (Table 1). The dataset aims to identify known and possible unknown BGCs placing them together with previously characterized PKSs from other fungi and bacteria. The genomes were downloaded using phylociraptor software, which automatically downloads genomes from NCBI and combines them with additionally specified genomes provided by the user [38]. We kept the original taxon names from NCBI for convenience, even though the current taxonomic status might differ.

**Table 1.** Genome basics for representatives of Ramalinaceae

| Nuclear genome | *Bacidia rubella* | *Bacidia gigantensis* | *Ramalina intermedia* | *Ramalina peruviana* |
| --- | --- | --- | --- | --- |
| Sequencing method | Illumina NovaSeq SP; MinION | PromethION 24 | Illumina MiSeq | Illumina MiSeq |
| Substrate | bark | bark | rock | twig |
| Growth form | crustose | crustose | fruticose | fruticose |
| Source | Axenic culture | Whole thallus | Whole thallus | Axenic culture |
| Raw reads produced | 17 Gbp; 22 Gbp (roughly) | 32 Gbp | 13.33 Gbp | - |
| Coverage | 500x | 500x | 290x | 2.0x |
| Assembly size (Mb) | 33.52 | 33.11 | 26.19 | 25.53 |
| Largest scaffold (bp) | 2.353.056 | 3.530.911 | 898.913 | 694.821 |
| Average scaffold (bp) | 657.358 | 1.379.912 | 148.821 | 25.764 |
| Number of scaffolds | 51 | 24 | 176 | 991 |
| N50 | 1.771.855 | 1.807.239 | 282.362 | 43.940 |
| GC (%) | 45.28 | 44.67 | 51.90 | 50.58 |
| Number of predicted genes | 8.773 | 8.451 | 7.405 | 6.756 |
| Number of predicted proteins | 8.728 | 8.400 | 7.355 | 6.706 |
| Number of unique proteins | 2.514 | 2.343 | 1.099 | 1.088 |
| tRNAs | 45 | 51 | 50 | 50 |
| **Proteins with at least one ortholog** | 5.860 | 5.711 | 6.111 | 5.467 |
| **Single-copy orthologs** | 2.345 | 2.345 | 2.345 | 2.345 |
| **Secreted proteins (SignalP)** | 678 | 612 | 464 | 384 |

### Genome annotation

The publically available genomes were sequenced with different sequencing technologies at different times and are thus of varying quality. To make annotations comparable, we performed *ab initio* gene calling and functional annotations for all of them. The downloaded genomes were analyzed using the smsi-funannotate (https://github.com/reslp/smsi-funannotate) pipeline based on funannotate v.1.8.7 [39]. First, we removed duplicated identical contigs (funannotate clean) in each assembly. To avoid long contig/scaffold names, we sorted our assembly by contig length and then renamed the fasta headers (funannotate sort). Afterwards, we made a soft repeat masking of the assembly using tantan (funannotate mask) [40]. The following steps included gene prediction using the gene-callers Augustus v3.3.2 [41], snap [42], GlimmerHMM v3.0.4 [43] and Genemark ES v4.68 [44]. For Augustus, we used *Aspergillus nidulans* as a pre-trained species. All the pipelines were run on the cluster of the University of Graz and our in-house Linux Server at LMU Munich. The genome statistics are summarised in Table 4. The output files of the newly annotated genomes from NCBI used in this study are available on figshare.

### Annotation of biosynthetic gene clusters

Identification, annotation and analysis of secondary metabolite BGCs in the studied fungal genome sequences were performed with antiSMASH v6.0 as a part of the smsi-funannotate pipeline. We chose all BGCs from the annotation results that contained orthologous core TI-PKSs for our comparative genomic and phylogenetic analyses. The GBK output files with annotated BGCs and predicted genes were visualized on the antiSMASH webserver (fungal version; accessed: November 2021) [45].

### Identification of atranorin biosynthetic gene cluster and phylogeny

To identify atranorin candidate genes, we downloaded sequences from PKS16 of *Cladonia grayi* (GenBank: ADM79459; 2089 aa) and PKS23 of *Stereocaulon alpinum* (GenBank: QXF68953; 2500 aa). Both PKSs were previously reported as possible candidates involved in the biosynthesis of atranorin (see Atranorin in Results and Discussion for details).

We used BLASTP v2.9.0+ [46] to detect orthologs of those characterized sequences in all PKS sequences we identified on the twenty-three studied genomes. We filtered BLAST output to retain sequences with a minimum identity of 30% over the alignment length and a minimum query coverage of 50%, sorted for the highest bit score and lowest e-value. Additionally, we used the best blast subject hits with the highest percent similarity to the query gene. The best hits from the blast results (>30%) were selected for the PKS23 alignment. The taxon selection was confirmed with the clade from the large TI-PKS phylogenetic tree.

We aligned the 29 identified amino acid sequences using MAFFT v7.480 [47] and calculated a maximum-likelihood using IQ-TREE v2.1.4 [48] with 1000 ultrafast bootstrap replicates, after selecting the best-fitting substitution model (LG+G8+F) with ModelFinder [49]. The final alignment comprised 29 sequences from 16 taxa, with 2320 amino acid sites, 2071 distinct patterns, 1756 parsimony-informative, 221 singleton sites, and 343 constant sites.

### Topology test to confirm atranorin biosynthetic genes

Our maximum-likelihood phylogeny recovered the *B. rubella* (Ramalinaceae) atranorin gene (bacrubpred_000804) as sister to a clade comprised of *Cladonia rangiferina* and *S. alpinum* (claranpred_005882 and QXF68953, respectively), which both belong to Cladoniaceae. Miadlikowska *et al*. [50] recovered Parmeliaceae as the closest relative of Cladoniaceae, with a clade of these two families being sister to Ramalinaceae. Thus, we tested if the monophyly of a clade comprising *Bacidia* + Cladoniaceae is significantly supported against the expected phylogenetic relationship comprising Parmeliaceae + Cladoniaceae. For this, we compared 1) an unconstrained ML tree to recover *Bacidia* + Cladoniaceae as monophyletic, and 2) a constrained tree with *Bacidia* sister to Cladoniaceae + Parmeliaceae (Fig. 1).

**Figure 1.**
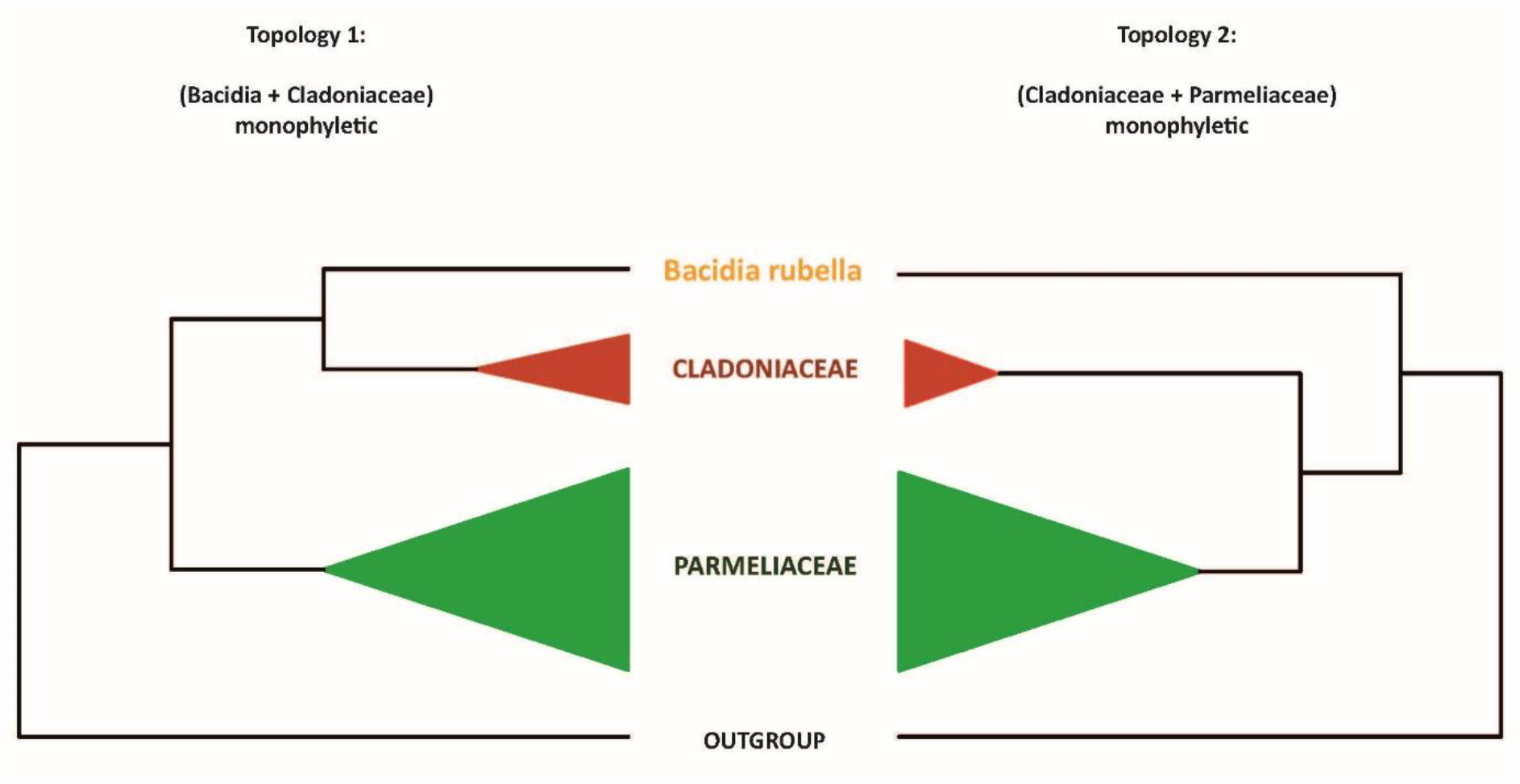
Two alternative topologies used for comparison in the topology test. Topology 1, recovering *Bacidia* + Cladoniaceae as monophyletic; Topology 2, recovering Cladoniaceae + Parmeliaceae as monophyletic.

For this topology test, we used PKS23 sequences in the strict sense, i.e., from the species known as atranorin produces (Table 2) with *Gyalolechia flavorubescens* and *Xanthoria elegans* as outgroups. The constrained tree can be multifurcating and need not contain all species; therefore, finally, we shortened our tree to the taxa we wanted to test. First, we performed a constrained search for both topologies using the LG model for the amino acid dataset. Then we concatenated both trees and implemented tree topology tests, based on the Kishino-Hasegawa test [51], Shimodaira-Hasegawa test [52], Expected Likelihood Weight [53], and approximately unbiased (AU) test [54] using IQ-TREE (-m LG -z concatenated_trees.treels -n 0 -zb 10000 -au). We considered tree topology to be unlikely if its test *p*-value was < 0.05 (marked with a “−” sign). The trees were visualized and examined using FigTree 1.3.1 [55].

**Table 2.** Overview of biosynthetic gene clusters and polyketide synthase families found in the twenty-three studied fungal genomes. Occurrence of the major secondary substance of *B. rubella*, atranorin, is highlighted. All sequences except *de-novo* sequenced *B. rubella* genome are from NCBI.

| Species | Species tag | Genome Size (Mb) | Number of Clusters | Type I PKS (total) | Type I NR-PKS | Type I R-PKS | PR-PKS | Hybrid PKS-NRPS | NRPS/putative NRPS | Metabolites reported |
| --- | --- | --- | --- | --- | --- | --- | --- | --- | --- | --- |
| *Alectoria sarmentosa* | alesarpred | 39.5 | 36 | 12 | 5 | 7 | 0 | 1 | 2/12 | Usnic acid, alectoronic acid (major), thamnolic, squamatic and barbatic acids |
| *Bacidia gigantensis* | bacgigpred | 33.1 | 31 | 11 | 7 | 4 | 0 | 1 | 5/10 | Homosekikaic acid |
| *Bacidia rubella* | bacrubpred | 33.5 | 31 | 10 | 6 | 4 | 0 | 0 | 3/7 | **Atranorin** |
| *Cladonia grayi* | clagrapred | 34.4 | 48 | 21 | 8 | 12 | 1 | 0 | 2/11 | 4-O-demethylgrayanic acid, colensoic acid, confumarprotocetraric acid, divaronic acid, fumarprotocetraric acid, grayanic acid, protocetraric acid, stenoporonic acid |
| *Cladonia macilenta* | clamacpred | 36.8 | 55 | 25 | 15 | 10 | 0 | 2 | 3/9 | Thamnolic acid, barbatic acid, didymic acid, squamatic acid, usnic acid, rhodocladonic acid |
| *Cladonia metacorallifera* | clametpred | 36.6 | 51 | 24 | 10 | 14 | 0 | 1 | 1/11 | Usnic acid, didymic acid, squamatic acid, rhodocladonic acid |
| *Cladonia rangiferina* | claranpred | 34.5 | 68 | 34 | 14 | 20 | 0 | 2 | 3/12 | **Atranorin**, protocetraric acid, fumarprotocetraric acid |
| *Cladonia uncialis* | clauncpred | 30.7 | 61 | 30 | 15 | 14 | 1 | 2 | 1/11 | Usnic acid, squamatic acid |
| *Cyanodermella asteris* | cyaastpred | 28.6 | 35 | 11 | 5 | 5 | 1 | 1 | 3/10 | Astin, skyrin |
| *Dibaeis baeomyces* | dibbaepred | 34.2 | 55 | 27 | 11 | 15 | 1 | 0 | 4/14 | Baeomycic acid, squamatic acid |
| *Endocarpon pusillum* | endpuspred | 36.2 | 32 | 14 | 4 | 9 | 1 | 1 | 3/4 | Not reported |
| *Evernia prunastri* | eveprupred | 40.2 | 86 | 36 | 16 | 19 | 1 | 4 | 3/20 | Usnic acid, **atranorin**, and chloroatranorin, evernic acid |
| *Graphis scripta* | grascrpred | 34.8 | 54 | 21 | 6 | 15 | 0 | 1 | 6/14 | Not reported |
| *Gyalolechia flavorubescens* | gyaflapred | 34.4 | 41 | 16 | 8 | 8 | 0 | 1 | 3/8 | Parietin, emodin, fallacinal, fragilin |
| *Lasallia hispanica* | lashispred | 39.7 | 28 | 15 | 8 | 6 | 1 | 1 | 0/3 | Gyrophoric acid, lecanoric acid, umbilicaric acid, skyrin |
| *Lasallia pustulata* | laspuspred | 32.9 | 26 | 17 | 9 | 7 | 1 | 0 | 0/4 | Gyrophoric acid, lecanoric acid, hiascinic acid, skyrin |
| *Letharia columbiana* | letcolpred | 52.2 | 43 | 14 | 7 | 7 | 0 | 2 | 3/6 | Vulpinic acid and **atranorin** |
| *Letharia lupina* | letluppred | 49.2 | 48 | 18 | 11 | 7 | 0 | 2 | 3/11 | Vulpinic acid and **atranorin** in the cortex, with norstictic acid in the hymenium of the apothecia |
| *Pseudevernia furfuracea* | psefurpred | 37.8 | 48 | 26 | 8 | 18 | 0 | 3 | 3/11 | **Atranorin**, physodic acid and oxyphysodic acid |
| *Ramalina intermedia* | ramintpred | 26.2 | 54 | 31 | 13 | 17 | 1 | 3 | 5/9 | Usnic acid (major); medulla with homosekikaic acid (major), sekikaic acid (major), 4'-O-methylnorhomosekikaic acid (minor) |
| *Ramalina peruviana* | ramperpred | 25.5 | 43 | 17 | 9 | 7 | 1 | 1 | 4/10 | Usnic acid, homosekikaic acid (major), sekikaic acid (major), and 4'-O-methylnorhomosekikaic acid and 4'-O-methylnorsekikaic acid (minor) |
| *Usnea hakonensis* | usnhakpred | 40.4 | 70 | 23 | 10 | 12 | 1 | 3 | 5/22 | Usnic and norstictic acids |
| *Xanthoria elegans* | xanelepred | 44.2 | 63 | 25 | 7 | 18 | 0 | 1 | 7/17 | Parietin (major), fallacinal, emodin, teloschistin and parietinic acid |

### Identification of orthologues and orthogroups with BUSCO and OrthoFinder

Orthofinder uses a hybrid approach based on sequence similarity estimated by a blast all-vs-all search with diamond and subsequent reconstruction of gene trees and a species tree to identify (single copy) orthologs and paralogs. To identify orthologues of extracted PKSs, we inferred orthogroups with Orthofinder v2.5.4 [56] for 1) all predicted proteins and 2) all extracted TI-PKSs, independently. The independent runs for two datasets were conducted to check for the concordance of the final results. As the results did not contradict, we discussed the result of TI-PKS only. The Markov Cluster (MCL) inflation parameter was set up by default (1.5). The trees utilized by Orthofinder were reconstructed using Fasttree v2.1.10 (-M msa, -A muscle, -S diamond (default)) [57].

### Type I Iterative PKS alignment

We performed a phylogenetic analysis of all TI-PKSs identified in twenty-three studied genomes of Lecanoromycetes, to assess the relationship of non-reducing PKSs (NR-PKS) and reducing PKSs (R-PKS) (including partly reducing PKSs). First, we prepared a list of all identified TI-PKS based on our functional annotations. We retrieved the sequences based on the corresponding gene ID numbers included in the gff3 file of each genome using a custom python script (select_transcript.py: https://github.com/reslp/genomics/blob/master/select_transcripts.py; accessed May 2021). We included predicted ketoacyl synthase (KS) amino-acid alignment reported by Kroken *et al*. [14], to enhance our dataset. The inclusion of reference PKSs enables us to compare the tree topology proposed in Kroken *et al*. [14] to our larger sampling of putative PKS genes.

All TI-PKS sequences were combined into a single file and aligned using MAFFT v7.480 [47]. We chose the E-INS-i alignment strategy because it performs better when aligning sequences with several conserved motifs interspersed in long, unalignable regions. We trimmed the alignment using trimAl v.1.4.rev15 [58]. Initial testing of different trimming settings (e.g., - gappyout, -strict, and -automated), showed that parameter combination (-gt 0.70 -resoverlap 0.70 -seqoverlap 60) is a good trade-off between removing ambiguously aligned sites and keeping phylogenetically informative sites of the PKS sequences. The final data matrix consisted of 624 amino acid sequences with 1049 amino acid sites, 1049 distinct patterns, and 1043 parsimony-informative sites from 61 taxa (23 studied genomes and 38 other fungal and bacterial taxa from Kroken *et al*. [14]). In addition, PKS23 from *Stereocaulon alpinum* (QXF68953) and PKS16 from *Cladonia grayi* (ADM79459) from NCBI were included. The alignment is available on figshare.

We selected the best-fitting substitution model (LG+F+G4) according to the Akaike and Bayesian Information Criteria using the ModelFinder implemented in IQ-TREE [49] on our TI-PKS alignment. We calculated a phylogenetic tree using maximum likelihood (ML) analysis implemented in IQ-TREE v2.1.4 [48], with 1000 ultrafast bootstrap replicates [59]. We also calculated a tree using RAxML-NG v1.0.3 [60] with parameters --all --bs-trees 100 --model LG+F+G4 --threads 16 --data-type AA. The inferred phylogenetic tree was then rooted with Bacterial Type II polyketide sequences using phyx [61].

To compare the congruence of the trees, we utilized Dendroscope v3.7.6 [62] using Tanglegram (Algorithms). The resulting tree was visualized using FigTree 1.3.1 [55] and a custom R script, with additional annotations added in Adobe Illustrator v24.0.3.

## Results and Discussion

### I. General characteristics of *de-novo Bacidia rubella* genome

We report the *de-novo* assembled genome of the lichen-forming fungus *Bacidia rubella* obtained from an axenic fungal culture. The genome was sequenced using Illumina short-read combined with Oxford Nanopore long-read technology. This hybrid approach resulted in a high-quality genome assembly of *B. rubella*. The final assembly had a size of 33.52 Mb in 51 scaffolds, an N50 of 1.77 Mb (Table 1), and was 98% BUSCO complete (fungi_odb9). Using multiple *ab-initio* gene-calling methods, 8773 genes were identified (Table 1). The similarity of standard genome metrics between *B. rubella* and *B. gigantensis* [63] suggests a high quality of the *B. rubella* genome provided here (Tables 1).

The genome of *B. rubella* contains 31 BGCs (Table 2), including six non-reducing and four reducing TI-PKS sequences. This TI-PKS biosynthetic arsenal is similar to that of *B. gigantensis*, containing 11 identified TI-PKS genes (seven NR-PKSs and four R-PKSs, respectively; Table 2). Despite the overall similarity of TI-PKS gene numbers between *B. rubella* and *B. gigantensis*, our BLAST-based comparison of these biosynthetic clusters showed only two significant hits between the two species, one R-PKS (bacgigpred_003963 and bacrubpred_000636) and one NR-PKS (bacgigpred_008278 and bacrubpred_000202), with 62.8 and 63% similarity of the amino-acid sequences, respectively. This result is not surprising, given that both fungal species differ in their secondary metabolite profiles. The only secondary compound of *B. rubella* is atranorin, the most widespread secondary metabolite found in the genus *Bacidia* s. lat. (Ekman 1996) [64]. In contrast, *B. gigantensis* is currently the only known *Bacidia* species to produce homosekikaic acid [65].

The two species *Ramalina intermedia* and *R. peruviana* belong to the same family as the genus *Bacidia*, but differ in having many more BGCs as compared to the two *Bacidia* species. In detail, *R. intermedia* contains 54 BGCs, including thirteen non-reducing, seventeen reducing and one partially reducing TI-PKS sequences. *Ramalina peruviana*, on the other hand, contains 43 BGCs, including nine non-reducing, seven reducing and one partially reducing genes respectively; Table 2).

The two species *Ramalina intermedia* and *R. peruviana* belong to the same family as the genus *Bacidia*, but differ in having many more BGCs as compared to the two *Bacidia* species. In detail, *R. intermedia* contains 54 BGCs, including thirteen non-reducing, seventeen reducing and one partially reducing TI-PKS sequences. *Ramalina peruviana*, on the other hand, contains 43 BGCs, including nine non-reducing, seven reducing and one partially reducing genes respectively; Table 2).

#### Atranorin

Atranorin is the major secondary compound known in *B. rubella* [64] and received attention in previous studies on different lichen-forming fungi. PKS16 genes in *Cladonia rangiferina* [26] and PKS23 genes in four *Cladonia* species and *Stereocaulon alpinum* [29] have been proposed to be involved in atranorin biosynthesis. In the framework of our phylogenetic results including these previously identified sequences, a putative PKS23 gene involved in atranorin production could be identified in *B. rubella*. Our results are consistent with Kim *et al*. [29] grouping PKS23 sequences together with sequences from atranorin-producing lichen-forming fungi in the TI-PKS phylogeny (Fig. 2). The newly identified, putative PKS23 sequence of *B. rubella* was recovered as sister to PKS23 sequences from *C. rangiferina* and *S. alpinum*, suggesting a sister-group relationship of *Bacidia* with Cladoniaceae, whereby the PKS23 sequences from Parmeliaceae formed a sister-group to that clade (Fig. 2).

**Figure 2.**
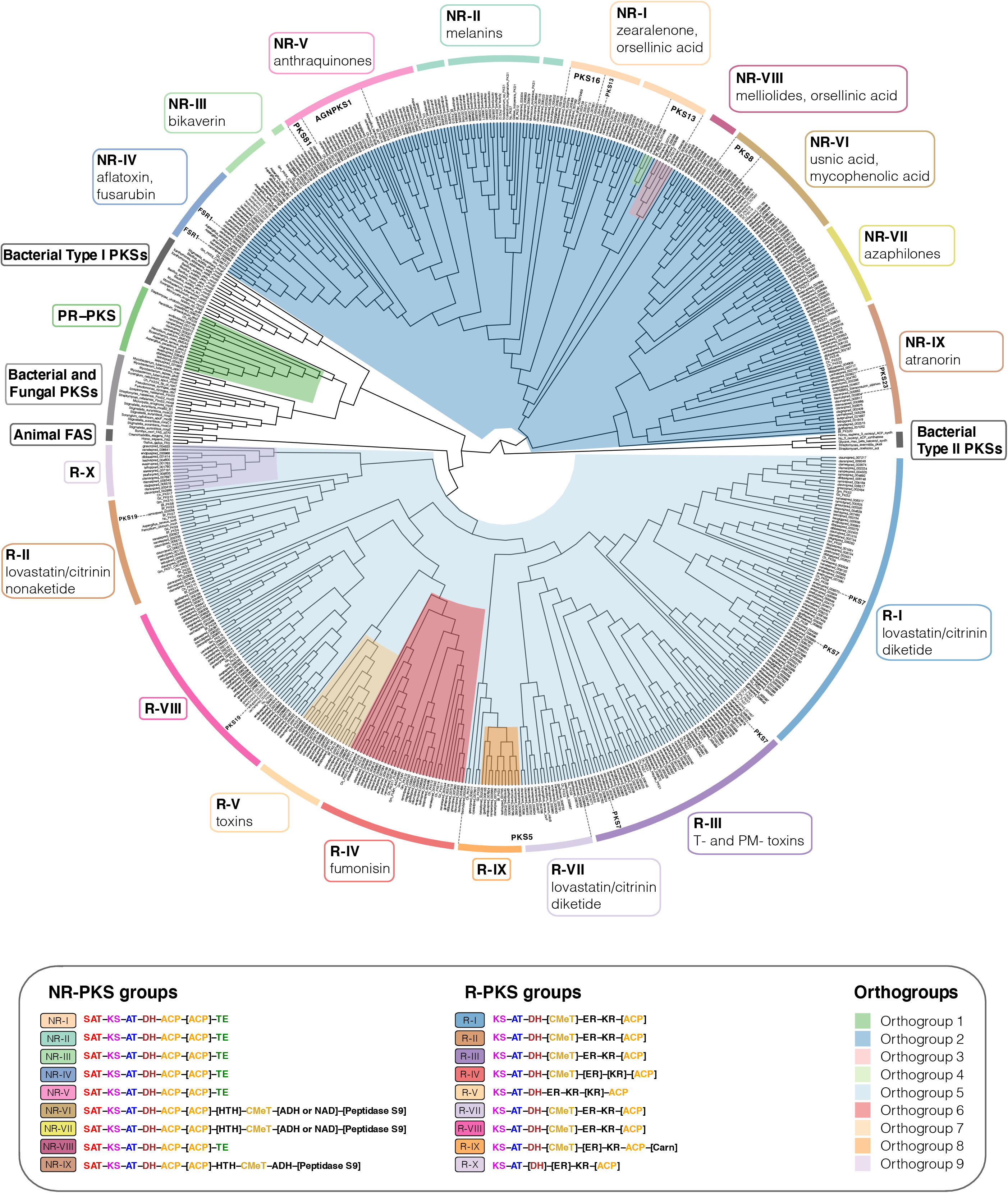
Maximum-likelihood phylogeny of Type I PKS genes inferred by IQ-TREE using Type II Bacterial PKSs as outgroup. All clades containing lichen-forming fungi are highlighted and the corresponding Orthogroups (1 to 9, respectively) are indicated by colour. The PR-PKS group corresponds to Orthogroup 1 (green); the nine NR-PKS groups (NR-I to NR-IX) belong to Orthogroups 2, 3 and 4 (dark-blue, pink and light-green, respectively); the ten R-PKS groups (R-I to R-X) belong to Orthogroups 5, 6, 7, 8, and 9 (light-blue, red, peach, orange, and lilac, respectively). Characteristic secondary substances for the groupings are given in the corresponding coloured boxes. Groups not containing lichen-forming fungal genes are indicated by grey boxes. For each group the domain arrangement of PKS is highlighted with distinct colors: SAT - starter unit-ACP transacylase, KS - ketoacyl synthase, AT - acyltransferase, ACP - acyl carrier protein, KR - ketoreductase, DH - dehydratase, ER - enoyl reductase, CMeT - methyltransferase, and TE - thioesterase, HTH - helix-to-helix, ADH - adhydrolase, NAD - NAD-binding, Carn - Choline/Carnitine O-acyltransferase domain.

This contrasts family-level taxonomic results with respect to morphology and phylogeny as detailed, e.g., by Peršoh *et al*. [66] and Miadlikowska *et al*. [50], where Cladoniaceae are more closely related to Parmeliaceae than to Ramalinaceae. In order to test the monophyly of the clade comprising *Bacidia* + Cladoniaceae compared to a group comprised of Parmeliaceae and Cladoniaceae, we performed a phylogenetic tree topology test on the PKS23 dataset. Specifically, we compared the unconstrained ML tree recovering *Bacidia* + Cladoniaceae as monophyletic (Topology 1) versus the constrained tree recovering Cladoniaceae + Parmeliaceae as monophyletic and *Bacidia* sister to that branch (Topology 2; Fig. 1).

Our results showed that the topology constraining Cladoniaceae + Parmeliaceae as monophyletic is significantly less likely (0.8%, Fig. 1, Table 3). One possible explanation is that the ancestor of *Bacidia* may have acquired the PKS23 gene from an ancestor of Cladoniaceae via horizontal gene transfer. It is clear that this hypothesis requires additional testing and an extended sample of PKS23 sequences from Cladoniaceae, Parmeliaceae, and Ramalinaceae. If additional studies do confirm this, it may also explain the scattered occurrence of atranorin in Ramalinaceae.

**Table 3.** Parameters for the topology test. Topology 1, recovering *Bacidia* + Cladoniaceae as monophyletic; Topology 2, recovering Cladoniaceae + Parmeliaceae as monophyletic. Value for the Kishino-Hasegawa test (Kishino and Hasegawa, 1989), Shimodaira and Hasegawa, 1989), expected likelihood weights (Strimmer and Rambaut, 2002) and approximately unbiased (AU) test (Shimodaira, 2002) are given.

| Tree | logL | deltaL | p-KH | p-SH | c-ELW | p-AU |
| --- | --- | --- | --- | --- | --- | --- |
| **Bacidia + Cladoniaceae** | -28054.52811 | 2.062e-08 | 0.495 + | 0.814 + | 0.496 + | 0.566 + |
| **Cladoniaceae + Parmeliaceae** | -28108.54142 | 54.013 | 0.0126 - | 0.0126 - | 0.00853 - | 0.0081 - |

deltaL: logL difference from the maximal logl in the set.
p-KH: p-value of one sided Kishino-Hasegawa test (1989).
p-SH: p-value of Shimodaira-Hasegawa test (2000).
c-ELW: Expected Likelihood Weight (Strimmer & Rambaut 2002).
p-AU: p-value of approximately unbiased (AU) test (Shimodaira, 2002).

Even though, PKS16 was first proposed to be involved in grayanic acid production in *Cladonia grayi* (CgPKS16) [18], it was later proposed to be involved in atranorin production in *C. rangiferina* as well [26]. In our phylogeny, these genes were placed together with several genes from other *Cladonia* and lichen-forming fungi (PKS16 clade, Fig. 2), but not all of them are known as atranorin producers (Table 2). Moreover, the contribution of PKS16 to atranorin production stays unclear, as atranorin is a B-orcinol depside, and therefore, the PKS producing it must have a CMeT domain for adding a methyl group to a carbon of the depside ring. But this is not the case for PKS16 thought to be involved in the synthesis of orcinol depsides [25]. The comparison of known metabolites in lichen-forming fungi containing PKS16 does not reveal any correlation and clear conclusion (Fig. 2, Group NR-I, PKS16 clade; Table 2). This indicates that the biosynthetic role of PKS16 remains elusive, and it seems to be involved in the biosynthesis of several secondary metabolites or a substance that has not been identified for these lichen-forming fungi yet.

#### Homosekikaic acid

Homosekikaic acid is a major secondary metabolite in *Bacidia gigantensis* as well as in both *Ramalina intermedia* and *R. peruviana*. However, it has not been found in *B. rubella*. To our knowledge, there was no putative PKS reported as being involved in the biosynthesis of homosekikaic acid and its biosynthesis has not been characterized using gene expression or heterologous expression, either. To identify candidate genes involved in the biosynthesis of homosekikaic acid in these three species, we used BLASTp on all predicted BGC sequences from the two *Bacidia* and *Ramalina* species. We identified three BGC candidates with blast similarity of the sequences ranging from 53 to 93%, but none of them was a TI-PKS. Instead, we recovered Type 3-PKS homologs from *B. gigantensis* (BGC 1.2), *R. intermedia* (BGC 6.1), and *R. peruviana* (BGC 223.1) as potential homosekikaic biosynthetic genes, but none of *B. rubella* BGCs showed high percent similarity to them. In addition, our TI-PKS phylogeny did not reveal any clade with sequences from both *Ramalina* species and *B. gigantensis*, but excluding *B. rubella*. One explanation is that the BGC responsible for the production of homosekikaic acid is still present in *B. rubella*, but the gene is not expressed, and thus, homosekikaic acid is not produced. Additional evidence from gene expression and analyses of substance profiles will be necessary to test this hypothesis. At this point, the genes involved in homosekikaic production remain to be found.

#### *Other biosynthetic genes identified* in silico *in the two* Bacidia *species*

Additional *in-silico* analyses using antiSMASH identified genes encoding enzymes to synthesize secondary metabolites included clavaric acid (100% similarity) and squalestatin S1 (40% similarity). Both terpenes were identified in both *Bacidia* species. A possible monascorubrin biosynthetic gene has been identified in *B. rubella*, showing 100% blast identity with the monascorubrin biosynthetic gene from *Talaromyces* (*Penicillium*) *marneffei* (PKS3: HM070047) [67]. Besides monascorubrin biosynthesis, PKS3 is also involved in the production of a well-known toxin, citrinin, as well as a yellow pigment, ankaflavin [67]. Monascorubrin and its related compounds are polyketides used as natural red colourants for a wide range of food [68]. In *B. rubella*, monascorubrin could be responsible for the characteristic orange to the orange-brown colouration of the apothecia. However, none of these substances has been reported for *B. rubella*.

In the genome of *B. gigantensis*, we identified genes highly similar to genes involved in the production of pyranonigrin E (100% similarity), naphthopyrone (100% similarity), and melanin (100% similarity). However, in most cases, only a part of the sequences showed high percent similarity; therefore, our *in-silico* based report is provisional, and detailed additional studies are necessary for unambiguous functional assignments.

### II. Biosynthetic gene composition in twenty-three annotated fungal genomes

The basic statistics for all twenty-three studied genomes are provided in Table 4. According to BUSCO homology searches against the fungal dataset (fungi_odb9), most genomes were of high completeness. We included only two genomes with completeness below 90%, namely *Alectoria sarmentosa* (75.8%) and *Graphis scripta* (88.6%). The number of predicted genes was in the range of 6756 to 11072. The lowest number of predicted genes was observed for *Ramalina peruviana* (most likely due to issues in the quality of the assembly given the lower sequencing depth), and the highest number was observed for *Evernia prunastri* (Table 4).

**Table 4.**
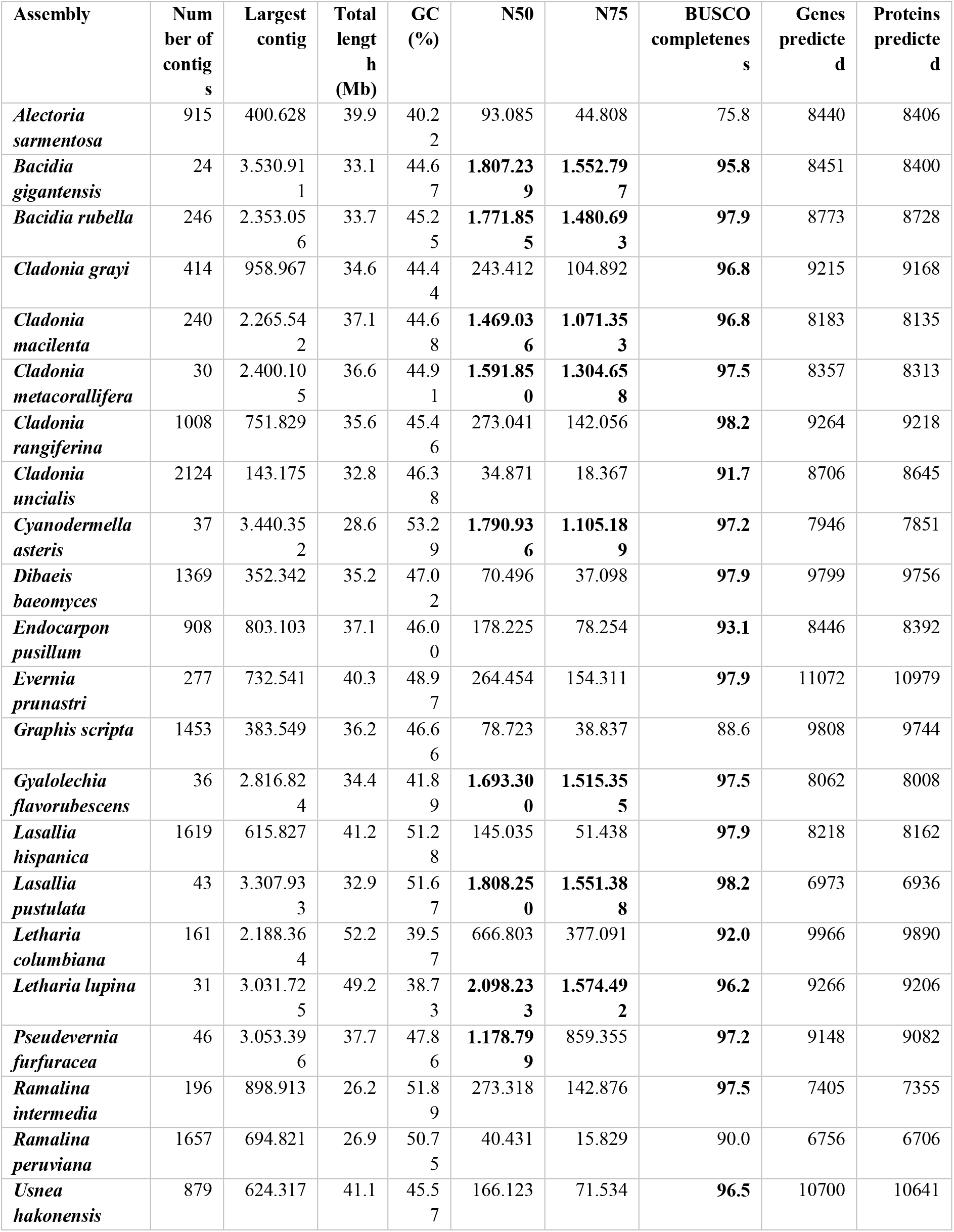
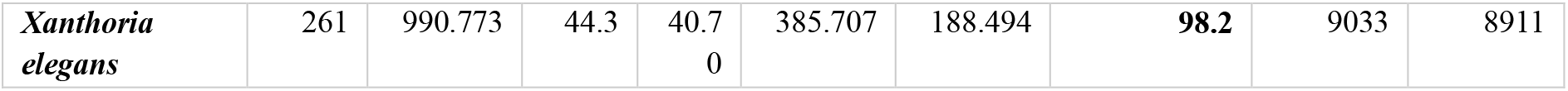
Genome basics for twenty-three studied fungal genomes

We investigated the BGCs predicted in all twenty-three studied genomes, belonging to different taxonomic groups and known to synthesize a large variety of secondary metabolites (Table 2 & 4). Our results revealed a high number of BGCs, with an average of 48 clusters per genome. The smallest number was recovered in the genome of *Lasallia pustulata* (26 BGCs), and the highest in *Evernia prunastri* (86 BGCs). In nearly half of the genomes, NR-PKSs are more numerous than R-PKS (Table 2). This contrasts with previous results reporting R-PKS gene numbers exceeding the number of NR-PKS genes [6]; therefore, a more extensive data set is necessary to evaluate if R-PKS or NR-PKSs are more numerous in the genomes of lichen-forming fungi.

The total number of TI-PKSs identified in all studied genomes was 478. Of those, 44.35% were of the non-reducing (NR), 53.35% of reducing (R), and 2.3% of partly reducing (PR) PKSs. The highest number of PKS genes was found in *E. prunastri* (36 TI-PKS clusters), and *C. rangiferina* (34), and the lowest numbers were in *B. rubella* (10), *B. gigantensis* (11), and *Cyanodermella asteris* (10) (Table 2).

Our extended genomic sampling also provides additional phylogenetic context for the NR-PKS genes of *B. rubella* and *B. gigantensis*. The six TI-PKS genes from *B. rubella* are from NR-PKS (three in Subclade I and three in Subclade II), while for *B. gigantensis*, seven TI-PKS genes are from NR-PKS (all are in Subclade I).

The discrepancy between the high number of recovered TI-PKS sequences and the small number of experimentally verified secondary metabolites (Table 2) raises the question about the role these genes play in the secondary metabolism and the products they produce. However, results linking PKS genes to lichen secondary metabolites beyond *in-silico* methods are still scarce.

### III. Type I PKS phylogeny

Our maximum-likelihood phylogeny of TI-PKS with a total of 624 sequences recovered from twenty-three fungal genomes and enhanced with previously published sequences is the largest analysis of biosynthetic gene content of various TI-PKS genes in lichen-forming fungi to date.

The TI-PKSs phylogenetic tree is divided into six main subgroups (Fig. 2): Bacterial Type II PKS (including bacterial and mitochondrial ketoacyl-ACP-synthetases; used as outgroup), Bacterial Type I PKS (also including some fungal sequences), Animal fatty acid synthase (FAS), PR-PKS, NR-PKS, and R-PKS. Lichen PKS genes were found in three of these main subgroups, viz. PR-PKS, NR-PKS, and R-PKS.

#### TI-PKS domain content mostly corresponds to phylogeny

TI-PKS genes encode for multi-domain enzymes, with each domain executing a defined function. The order and domain content of the PKSs thus defines the class of polyketides produced by the corresponding BGC. The presence or absence of particular domains has led to the classification of fungal PKSs into three main subgroups. The first subgroup comprises the NR-PKSs which do not contain reductive domains. The generalized domain content of NR-PKS in studied fungi in our phylogeny is SAT-KS-AT-DH-ACP-[ACP]-HTH-CMeT-[TE or ADH or NAD]-[Peptidase S9] (NR-I to NR-IX; Fig. 2). The second subgroup comprises PR-PKSs containing a single KR domain or KR and DH domains. Thus, the generalized domain content in PR-PKS in our phylogeny is KS-AT-[DH]-KR-ACP (PR-PKS; Fig. 2). Finally, R-PKSs contain a complete set of reductive domains, viz. KR, DH, and ER and thus exhibit the following domain structure: KS-AT-DH-CMeT-ER-KR-ACP-[Carn] (R-I to R-X; Fig. 2). The additional domains, such as helix-to-helix (HTH), adhydrolase (ADH), NAD-binding (NAD), Peptidase S9, and Choline/Carnitine O-acyltransferase domain (Carn) were not shown in the previous studies in the context of TI-PKS in lichen-forming fungi and are discussed in detail below in the corresponding sections.

#### The discrepancy of grouping PKS genes

Assigning PKS genes to different groups in fungi on the account of gene domain composition and their synthesized products were introduced by Kroken *et al*. [14] and later refined based on DH-domain pocket sizes by Ahuja *et al*. [69] and Liu *et al*. [70]. This classification was used by several studies on the investigation of PKS gene diversity in lichen-forming fungi, e.g., [29,71]. Although, it remains unclear if sequences from these groups indeed form monophyletic clades and if all genes from one group synthesize the same (or at least chemically similar) substances. In our phylogeny, previously proposed groups did not always form monophyletic groups (e.g., NR-II; Fig. 2) or characterized genes from one group have been shown to produce different substances (e.g., PKS81 and AGNPKS1 in NR-V; Fig. 2). To provide an additional and unbiased way to identify groups of sequences, we performed an Orthofinder run on all TI-PKS sequences from studied fungi. We identified 9 orthogroups named Orthogroup 1 to Orthogroup 9, respectively. Orthogroup 1 corresponds to the PR-PKSs (Fig. 2). Three orthogroups contain NR-PKSs (Orthogroups 2, 3, and 4), but none of them correspond explicitly to any of the nine groups identified before by Pizarro *et al*. [71] and Kim *et al*. [29]. Orthogroups 5, 6, 7, 8, and 9 contain R-PKSs, only two of which correspond to one of the eight groups defined in Kroken *et al*. [14] and Punya *et al*. [72]. Orthogroup 6 corresponds to R-IV, and Orthogroup 7 corresponds to R-V. All Orthogroups identified here correspond to supported clades in the PKS phylogeny (Fig. 2). The discrepancy between our results and previous studies will have to be investigated in subsequent studies. Nevertheless, several groups of the NR-PKS and R-PKS showed differences in the PKS domains present; those groups do contain support from different sources and thus merit discussion (see below).

#### Lichen-forming fungi contain only a few partially reducing PKSs in the phylogenetic neighbourhood to bacterial PKSs

The PR-PKS sequences formed a well-supported clade sister to bacterial Type I PKS sequences with other fungal PKSs (PR-PKS; Fig. 2). Their domain configuration is KS-AT-[DH]-KR-ACP, including only a single reductase domain (KR). In contrast to the large NR- and R-PKS groups, PR-PKS contains mainly genes from *Aspergillus* and *Penicillium* and only a few genes from lichen-forming fungi. Genes from the studied fungal genomes have the typical PR-PKS domain composition, but in two genes from two *Ramalina* (ramintpred_001715 and ramperpred_002012), the DH domain was missing. PR-PKS genes form a sister clade to bacterial genes only (Bacterial Type I PKSs; Fig 2). Bacterial T1-PKSs often possess a KR domain catalyzing the first step in the reductive modification of the beta-carbonyl centres in the growing polyketide chain. This domain requires NADPH to reduce the keto- to a hydroxy group [73].

The “simple” domain configuration of Bacterial and PR-PKS genes compared to NR- and R-PKSs has not escaped our attention. Our study was not designed to specifically test how PKS genes in Ascomycetes were acquired and how they diversified. However, the placement of Bacterial and NR-PKS sequences as the earliest branches in our phylogeny also supports the hypothesis suggested by Kroken *et al*. [14] that fungal TI-PKS genes could have been acquired by an ancient horizontal gene transfer between bacteria and fungi. Moreover, our phylogenetic results suggest that ancestral TI-PKS genes may have been partially reducing. Under this scenario, NR- and R-PKS evolution could have been connected with domain structure modification (such as gain of SAT in NR-PKS) and subsequent functional diversification. However, this cannot be concluded with certainty without a greatly expanded sample of genomes from other fungal groups, including genomes from early-branching fungal lineages and comprehensive ancestral state reconstructions.

### Fungal PKSs producing non-reduced polyketides

#### The diversity of NR-PKS sequences

In previous studies, NR-PKSs have been divided into nine major groups based on protein sequence similarity and PKS domain content [14,29,69,71]. Similar to previous studies, we observed characteristic domain configurations for the different clades in our phylogeny. The domain structure of NR-PKSs in studied fungi can be generalized as follows: SAT-KS-AT-DH-ACP-[ACP]-HTH-CMeT-[TE or ADH or NAD]-[Peptidase S9], however, some PKS genes may deviate from this structure. Observed domain content variations are not random but rather occur in two main patterns: either through duplication of the ACP domain or through the addition of an N-terminal TE domain. PKS genes coding for an additional ACP domain are scattered throughout different NR-PKS clades, suggesting multiple independent gains. However, the functional significance of ACP domain duplications is still unknown [14]. On the other hand, the TE domain is involved in a thioesterase-mediated product release which is the most common release mechanism in TI-PKS [74]. It regularly extends to a C-C Claisen cyclization domain (TE/CLC domain), e.g., in *Aspergillus parasiticus* PksA [75]. Only sequences from groups NR-I and NR-II have TE domains in our phylogeny. Although CYC domains have previously been reported from various fungi [14,69,70], we could not identify them in our analyses and thus did not indicate them in the phylogeny.

Most of the NR-PKS sequences belong to a large clade containing groups NR-I to NR-V and NR-VIII with domain configuration containing a TE domain at the N-terminal end.

#### Subclade I of NR-PKSs (including groups NR-I to NR-V and NR-VIII)

##### Group NR-I

The paraphyletic group NR-I contains several clades with previously characterized sequences involved in the biosynthesis of aromatic compounds derived from orsellinic acid, such as grayanic acid (PKS16), physodic acid and olivetoric acid [24,25], lecanoric acid [76], xanthones, aflatoxin, and naphthoquinones [14]. In addition, the clade also comprises PKS13 sequences from several lichen-forming fungi and *Gibberella zeae* as well as a PKS15 sequence from *Botryotinia fuckeliana* with unknown functions.

##### Group NR-II

Our results show that group NR-II is diverse, comprised of several clades containing sequences from lichen-forming fungi interspersed with characterized PKS genes proposed to be involved in the biosynthesis of different substances. The earliest branching clade in NR-II contains genes involved in the 6-hydroxymellein biosynthesis (NR-II, Fig. 2), a key intermediate in terrein biosynthesis [77]. Terrein, produced in large quantities by *Aspergillus terreus*, has phytotoxic activity and could potentially serve as a novel antibiotic [78]. Our phylogeny revealed sequences of four lichen-forming fungi (clagrapred_001463, clauncpred_002890, psefurpred_002946, and grascrpred_004317) clustering together with the *A. terreus* terA gene. An additional BLAST search of terA (GenBank: EAU38791) against these sequences revealed 58 to 67% similarity on the amino acid level, indicating a high degree of conservation despite an approximate 350 million years split of Eurotiomycetes (*Aspergillus*) and Lecanoromycetes [79]. Terrein has not been reported from lichen-forming fungi; therefore, without further experiments and detailed substance characterization, terrein production remains hypothetical.

Another important pattern is the occurrence of characterized melanin genes and melanin precursors in different clades. Melanins are a diverse group of substances that are synthesized via different pathways [80]. Melanins produced by PKSs are either 1,8-dihydroxynaphtalene (DHN) melanin or Deoxybostrycoidein-melanin which play a role in virulence, morphogenesis, or the response to environmental stress [81]. Melanins are insoluble and thus cannot be studied by standard biochemical methods [80]. In the PKS gene survey by Kroken *et al*. [14], all melanin synthesizing genes were grouped together in a single monophyletic clade. In our analysis, they are recovered in at least three clades containing sequences from several Lecanoromycetes and *Endocarpon pusillum*. Sister to NR-II is a clade containing sequences from *Colletotrichum lagenarium* (PKS1), *Glarea* sp., *Nodulisporium* sp., and *C. heterostrophus* (PKS18), which were also characterized as producers of melanin [14]. The high number of genes from lichen-forming fungi close to characterized melanin biosynthetic genes suggest an essential role of melanins in that group.

The crown clades in the NR-II group include the annotated PKS81 orthologs from *Cladonia metacorallifera*, *C. macilenta*, *Lasallia pustulata* and *L. hispanica* as sister to a PKS19 sequence from *Cochliobolus heterostrophus* and two clades with characterized Agnpks1 also from several lichen-forming fungi. Agnpks1 (Atrochrysone carboxylic acid synthase) was originally described from *Penicillium divaricatum* and is involved in the biosynthesis of agnestins and dihydroxy-xanthone metabolites [82]. Xanthones and related benzophenones are produced by various filamentous fungi. They exhibit insecticide, antioxidant, antibacterial, anti-inflammatory, and anticancer activities [83]. Examples include desmethylsterigmatocystin, a key intermediate of the aflatoxin group of mycotoxins produced by *Aspergillus flavus*, and a norlichexanthone from the lichen-forming *Lecanora straminea* [82]. An additional BLAST search of norlichexanthone synthase from *L. straminea* (GenBank: D7PI15) against our annotated fungal sequences showed at least 37% similarity to sequences from the clade containing PKS81 and Agnpks1. The relationship between these two genes remains unclear, but given our results, PKS81 might belong to or have evolved from Agnpks1. Additional closer investigations will be necessary to show the role of PKS81 and Agnpks1 orthologs in lichen-forming fungi and their possible role in xanthone biosynthesis.

##### Groups NR-III and NR-IV

Sister groups NR-III and NR-IV contain proteins synthesizing large polyketide chains, such as conidial yellow pigment Alb1 (NR-III) or aflatoxin/sterigmatocystin of *A. nidulans* (NR-IV) [14,71,84].

Group NR-III also contains characterized PKSs producing duclauxin (35-71% similarity) and naphthopyrone (100% similarity). Duclauxins are dimeric, heptacyclic fungal polyketides with various bioactivities [85], while bis-naphthopyrones act in herbivore defence in filamentous ascomycetes [86].

Several genes recovered in the NR-IV group are potentially involved in producing different fungal pigments. The genes from *Ramalina intermedia* (ramintpred_005964) and *Cladonia grayi* (clagrapred_006320) has been assigned to an FSR1 gene, involved in the production of the highly pigmented naphthoquinones fusarubins involved in fruiting body colouration of *Fusarium fujikuroi* [87]. An additional BLAST search with the *R. intermedia* gene (ramintpred_005964) showed 72.3% similarity to a PKS of *Cladonia metacorallifera* (GenBank: QIX11499), which is involved in the biosynthesis of the red compound cristazarin, having an antibacterial and antitumor activity [88]. *Bacidia rubella*, from which a sequence was recovered as sister to the putative *Ramalina intermedia* FSR1, and species of *Cladonia* exhibit red-pigmented fruiting bodies, while *R. intermedia* lacks red pigments [89]. This evidence raises the possibility that a combination of different PKS genes could produce red-coloured pigments (see also discussion on monascorubrin above) and are responsible for the red fruiting bodies in *B. rubella*.

##### Group NR-V

Group NR-V comprised PKSs without the TE domain, however in the characterized PKSs in our phylogeny, the TE or R domains were present in all clades. The genes in the NR-V group are suggested to be involved in the production of different mycotoxins, such as desertorin (*Aspergillus nidulans*) or atrochrysone (*Aspergillus fumigatus*) [71,90,91].

#### Subclade II of NR-PKSs (including groups NR-VI, NR-VII, and NR-IX)

The second large clade of NR-PKS sequences includes the previously recovered groups NR-VI, NR-VII, and the recently defined group NR-IX [29] (Fig. 2). PKSs in these groups also deviate from the general domain pattern mentioned above. Domain arrangement can be generalized as follows: SAT-KS-AT-DH-ACP-[ACP]-[HTH]-CMeT-[ADH/NAD]-[Peptidase S9]. Although TE domains were previously reported from all NR-PKS groups except group NR-V [69,71], we have not observed the presence of TE in groups NR-VI, NR-VII, and NR-IX. Instead, NAD and ADH domains were present, in some cases followed by a Peptidase S9 (PFAM: PF00326) domain. Peptidase S9 belongs to proteolytic enzymes consuming serine in their catalytic activity and they are ubiquitously found in viruses, bacteria, and eukaryotes [92]. We observed a helix-turn-helix domain (HTH) being inserted after one or two ACP domains in several groups. The HTH domain has been previously found in PKSs in fungi but it has not been shown in the arrangement of the NR-PKSs of lichen-forming fungi before. Proteins with an HTH motif are, for example, involved in DNA repair, RNA metabolism, and protein-protein interaction [93] and could thus be involved in developmental or morphogenic processes.

##### Group NR-VI

Characterized PKSs in this group include PKS18 and PKS19 genes from *Botryotinia fuckeliana*. It was suggested that genes of this group are involved in usnic acid biosynthesis [71] and belong to PKS8 according to Kim *et al*. [29]. Our results show that this group is divided into two clades. The first clade contains genes of known usnic acid-producers such as *Ramalina intermedia*, *R. peruviana*, *Cladonia metacorallifera*, *C. macilenta*, *C. uncialis*, *Evernia prunastri*, *Alectoria sarmentosa*, *Letharia columbiana*, and *Usnea hakonensis*, interspersed with genes from lichen-forming fungi known not to produce usnic acid. The second clade includes genes mostly from lichen-forming fungi not producing usnic acid, including a gene from *B. rubella* (bacrubpred_000551). However, sequences from *Cladonia macilenta*, *C. uncialis*, and *Evernia prunastri* were also recovered in this clade.

##### Group NR-VII

The only genes characterized in this group are PKS17 from *Botryotinia fuckeliana* and PKS21 from *Cochliobolus heterostrophus*. These genes have been proposed to produce citrinin or lovastatin [14,94,95]. The other identified genes from the lichen-forming fungi in this group had 50 to 70% similarity to monascorubrin, citrinin, conidial yellow, azanigerone A, and stipitatic acid (see also discussion on monascorubrin above). As mentioned above, the biosynthetic gene from *B. rubella* (bacrubpred_007642) has a 100% identity to the monascorubrin biosynthetic gene.

##### Group NR-IX

The recently recognized group NR-IX [29] contains several predicted PKS23 sequences, involved in the biosynthesis of atranorin (see sections on Atranorin and Homosekikaic acid above) and methylated orsellinic acid derivatives. In contrast to groups NR-VI and NR-VII, group NR-IX sequences always contain an ADH and lack a NAD domain after CMeT, while Peptidase S9 is only sporadically present. It is necessary to note that a single ACP domain was present in the PKS23, including atranorin producers. In addition, other characterized PKSs of this group are from *Cochliobolus heterostrophus* (PKS22 and PKS23) and *Botryotinia fuckeliana* (PKS16 and PKS20).

### Fungal PKSs producing reduced polyketides

The R-PKS genes are involved in synthesizing variously reduced, usually linear polyketides. They are often precursors of toxins that are active in animals (e.g., lovastatin, citrinin, and fumonisin) and of toxins that are active in plants (e.g., T-toxin and PM-toxin) [96–99]. We follow this classification in our discussion whenever possible, while also highlighting deviations. Compared to NR-PKSs, the possible roles of R-PKSs are less known. In our phylogeny, R-PKSs formed a clade of similar size and sister to NR-PKSs with the well-supported clades. Unlike what has been shown in previous studies, all studied R-PKSs here lacked an additional second ACP domain, and in some cases, ACP was not present. According to Kroken *et al*. [14], four major groups (R-I to R-IV) of R-PKS genes can be distinguished based on their overall domain structure and known compounds. This classification was later extended to groups R-V to R-VIII by Punya *et al*. [72]. Almost 72% of the studied fungal sequences could be assigned to groups designated by Kroken *et al*. [14] and Punya *et al*. [72] with the general domain content: KS-AT-DH-[CMeT]-ER-KR-[ACP]. Two clades could not be recognized from our tree. First is Clade VI, which included uncommon PKS-NRPS hybrids with four additional domains after ACP, viz. condensation (C), adenylation (A), thiolation (T), and reductase (R) [72]. But as our study focused on the TI-PKSs, it did not include hybrid NRPS-PKS genes. The second is Group VIII, whose sequences were not present in our phylogeny, and are thus not feasible to identify with confidence. In addition, based on our phylogenetic results, two R-PKS clades did not match to any of the previously reported R-PKS groups. Given the characteristic domain structure of these clades and because they were recovered as distinct orthogroups in our Orthofinder analysis, we propose those as two additional groups, R-IX and R-X, for these clades following the numbering scheme by Kroken *et al*. [14] and Punya *et al*. [72].

#### Subclade I of R-PKS (including R-I, R-III, R-VII, RV-III, and new R-IX and R-X groups)

##### Group R-I

Group R-I contains one of the R-PKS genes synthesizing the diketide portion of lovastatin and citrinin, T-toxin, and PM-toxin [14]. Our results show that the genes of this group, recovered in a monophyletic clade by Kroken *et al*. [14], are scattered throughout different clades in our analysis; therefore, R-I does not have a clear border.

##### Group R-II

Group R-II forms a well-supported clade and is characterized by the absence of an ER domain [14]. It includes many previously published sequences and orthologs from *Cladonia* and *Letharia* species, *Xanthoria elegans*, *Pseudevernia furfuracea*, and *Ramalina intermedia*. The *R. intermedia* ortholog was assigned to PKS19. PKS19 was predicted to be involved in the production of heptaketides with a role in a lesion of rice leaves formation in *Magnaporthe oryzae* [100]. Two characterized proteins from *A. terreus* lovB and *P. citrinum* mlcA synthesize the cyclic nonaketide portion of lovastatin and citrinin.

##### Group R-III

Group R-III forms a clade comprised of four sequences previously assigned to this group by Kroken *et al*. [14] and also contains a sequence from *Cyanodermella asteris*. PKSs from this clade have lost the ME domain. The only characterized gene in group R-III is *C. heterostrophus* PKS2, which, along with PKS1, is required for the synthesis of T-toxin [14]. We did not recover the genes of lichen-forming fungi in this group.

##### Group R-IV

The genes in the R-IV group may contain a conserved CMeT domain. Their domain configuration is thus similar to that of groups R-I and R-II. However, N-terminal domains ER, KR, and ACP may or may not be present in group R-IV. The only characterized PKS in this group is *G. moniliformis* FUM1 (Gm_FUM1_AAD43562), which makes the linear polyketide precursor of the toxin fumonisin [98]. This group contains several sequences from Kroken *et al*. [14] interspersed with sequences from various lichen-forming fungi.

##### Group R-V

Group R-V has similar enzymatic domains as Orthogroup 7 with the following domain configuration: KS-AT-DH-ER-KR-[KR]-ACP. The second KR domain is only present in *Dibaeis baeomyces* (dibbaepred_000831). Several lichen-forming fungi possess genes belonging to this group. Again, experimentally characterized genes from *Gibberella moniliformis* and *Botryotinia fuckeliana* indicate a possible role in toxin biosynthesis [14].

##### Group R-VII

This group contains several previously characterized sequences as well as sequences from *R. intermedia*, *P. furfuracea*, *G. furfuracea*, and *C. macilenta* with PKS5 annotations. Additional sequences from lichen-forming fungi had no functional annotations; therefore, the role of these genes in lichen-forming fungi remains unclear. The characterized sequences of *Aspergillus* and *Penicillium* indicate a role in the production of the diketide portion of lovastatin and citrinin but were assigned to group R-I by Kroken *et al*. [14].

##### Novel group R-IX

The newly defined group R-IX is sister to group R-VII and is characterized by an additional CMeT domain: KS-AT-DH-CMeT-ER-KR-ACP-[Carn]. It contains Orthogroup 8, and both groups differ from all other groups by having a facultative Choline/Carnitine O-acyltransferase domain (Carn) following the ACP domain: KS-AT-DH-ER-KR-[ACP]-[Carn] (Fig. 2). The Carn domain is found in several eukaryotic acetyltransferases. This domain is also found at the C terminus of a highly reducing polyketide synthase (SdnO), part of a gene cluster mediating the biosynthesis of glycoside antibiotics sordarin and hypoxysordarin [101].

##### Novel group R-X

The second novel group R-X, corresponding to Orthogroup 9, contains only genes from lichen-forming fungi. Together with Group R-II, they share a sister-group relationship to all other R-PKSs in our phylogeny (Fig. 2). Genes from this group may not have DH, ER, and ACP domains, resulting in the following domain content: KS-AT-[DH]-[ER]-KR-[ACP]. Most fungal genes recovered here are missing the DH and ER domain. However, the genes from *Lasallia hispanica* (lashispred_001760) and *Evernia prunastri* (eveprupred_001763) contain a complete domain configuration: KS-AT-DH-ER-KR-ACP.

### Conclusions & Limitations

This study presents a comprehensive analysis of the PKS gene content of the *de-novo* sequenced genome of *B. rubella* and twenty-two publicly available genomes of mainly lichen-forming fungi. Our results reveal a large diversity of PKS genes in lichen-forming fungi, much larger than expected based on their recorded secondary metabolite profiles. This highlights the potential of lichen-forming fungi to produce a much higher number of substances than previously assumed. Our *in-silico* approach provides first insights into the possible roles of these genes; however, we are aware that this comes with limitations. Only detailed studies of individual BGCs combining gene expression analyses with analytical chemistry and metabolomics will allow thorough testing of the hypotheses proposed here. Prospects for this are promising as an increasing number of studies succeeded in the heterologous expression of lichen-forming fungal genes, e.g., [29,76,88,102,103]. Subsequent *in vivo* studies of metabolite profiles in lichen-forming fungi supplemented by high-resolution MS/MS spectra [104] or metabolic profiling based on stable isotope analysis [105] will contribute to the knowledge on how secondary metabolites in lichen-forming fungi are produced. Results based on multiple experimental approaches that continue being incorporated into specialized databases such as antiSMASH will provide novel insights and help to formulate hypotheses about the biosynthetic potential of lichen-forming fungi, as the number of available lichen-forming fungal genomes continues to grow.

## Acknowledgments

We thank Dr. Diego F. Morales-Briones for his valuable help in the tree figure preparation. Andrea Brandl and Tanja Ernst helped with the lab work. JG was supported by a BAYHOST fellowship from the Bayerische Staatsministerium für Bildung und Kultus, Wissenschaft und Kunst and an LMU travel grant (April 2021). Laboratory work and genome sequencing were funded by the LMU Munich start-up funds of SW and by the Staatliche Naturwissenschaftliche Sammlungen Bayerns, grant SNSB innovativ to AB. We thank the government of the administrative district of Upper Bavaria for the sampling permit.

## Data Availability Statement

The data supporting the findings of this study will be available open access in figshare at http://doi.org/[xxx]. These include gff3 files and protein sequences from twenty-two re-annotated genomes downloaded from the NCBI. In addition, the alignment used for the PKS tree calculation will be deposited. The *de-novo* sequenced genome of *Bacidia rubella* will be available on the NCBI at link: xxx and No.

## Contributions

Conceptualization, J.G., A.B., S.W., and P.R.; methodology, software and formal analysis, J.G. and P.R.; investigation, J.G., and A.B.; resources and funding acquisition, S.W. and A.B.; data curation, J.G., and P.R.; writing—original draft preparation, J.G.; writing—review and editing, J.G., A.B., S.W., and P.R.; visualization, J.G.; supervision, A.B., S.W., and P.R. All authors have read and agreed to the published version of the manuscript.

